# Multiresolution functional brain parcellation in an elderly population with no or mild cognitive impairment

**DOI:** 10.1101/077974

**Authors:** Angela Tam, Christian Dansereau, AmanPreet Badhwar, Pierre Orban, Sylvie Belleville, Howard Chertkow, Alain Dagher, Alexandru Hanganu, Oury Monchi, Pedro Rosa-Neto, Amir Shmuel, John Breitner, Pierre Bellec, Alzheimer’s Disease Neuroimaging Initiative

**Affiliations:** McGill University, Montreal, QC, CA; Douglas Mental Health University I’nstitute, Research Centre, Montreal, QC, CA; Centre de recherche de l’institut universitaire de gériatrie de Montréal, QC, CA; Université de Montréal, QC, CA; University of Calgary, AB, CA; Hotchkiss Brain Institute, Calgary, AB, Canada

## Abstract

We present group brain parcellations for clusters generated from resting-state functional magnetic resonance images for 99 cognitively normal elderly persons and 129 patients with mild cognitive impairment, pooled from four independent datasets. The brain parcellations have been registered to both symmetric and asymmetric MNI brain templates and generated using a method called bootstrap analysis of stable clusters (BASC, Bellec et al., 2010). Eight resolutions of clusters were selected using a data-driven method called MSTEPS (Bellec, 2013). We present two variants of these parcellations. One variant contains bihemisphereic parcels (4, 6, 12, 22, 33, 65, 111, and 208 total parcels across eight resolutions). The second variant contains spatially connected regions of interest (ROIs) that span only one hemisphere (10, 17, 30, 51, 77, 199, and 322 total ROIs across eight resolutions). We also present maps illustrating functional connectivity differences between patients and controls for four regions of interest (superior medial frontal cortex, dorsomedial prefrontal cortex, striatum, middle temporal lobe). The brain parcels and associated statistical maps have been publicly released as 3D volumes, available in .mnc and .nii file formats on figshare (http://dx.doi.org/10.6084/m9.figshare.1480461) and on Neurovault (http://neurovault.org/collections/1003/). This dataset was generated as part of the following study: Tam A, Dansereau C, Badhwar A, Orban P, Belleville S, Chertkow H, Dagher A, Hanganu A, Monchi O, Rosa-Neto P, Shmuel A, Wang S, Breitner J, Bellec P for the Alzheimer’s Disease Neuroimaging Initiative (2015) Common Effects of Amnestic Mild Cognitive Impairment on Resting-State Connectivity Across Four Independent Studies. Front. Aging Neurosci. 7:242. doi: 10.3389/fnagi.2015.00242 Finally, the code used to generate this dataset is available on Github (https://github.com/SIMEXP/mcinet).

Specifications Table

|  |  |
| --- | --- |
| Subject area | *Biology* |
| More specific subject area | *Neuroscience* |
| Type of data | *Images (3D brain volumes)* |
| How data was acquired | *MRI, resting-state functional MRI (Philips & Siemens 3T scanners)* |
| Data format | *Analyzed* |
| Experimental factors | *Pre-processing for motion-related or other artifacts, group-level statistical analysis* |
| Experimental features | *We pooled resting-state fMRI data from 4 independent studies with cognitively normal elderly subjects and patients with mild cognitive impairment to generate 1) group-level functional brain parcellations with varying numbers of parcels, and 2) maps illustrating functional connectivity differences between patients and controls in four parcels of interest.* |
| Data source location | *Canada & The United States* |
| Data accessibility | Figshare: http://dx.doi.org/10.6084/m9.figshare.1480461<br>Neurovault: http://neurovault.org/collections/1003/<br>Github: https://github.com/SIMEXP/mcinet |

## Value of the data

- We have released group-level resting-state functional brain parcellations at multiple levels of spatial resolution derived from several samples of older adults with and without mild cognitive impairment (total of 228 subjects) and t-maps displaying functional connectivity differences between patients and controls for selected regions of interest, as well as the code used to generate these maps
- These parcellations have been selected to represent 8 different spatial resolutions, ranging from 4 to 208 bihemispheric clusters and 10 to 322 regions of interest.
- These parcellations can be used as atlases for brain imaging studies in elderly populations.
- The functional clusters and t-maps we have derived can be used as target regions in hypothesis-driven studies, especially for those interested in aging, mild cognitive impairment and dementia
- The code can be adapted to generate similar atlases on other datasets or populations

## Data

This work is derived from the Alzheimer’s Disease Neuroimaging Initiative 2 (ADNI2) and three samples from Montreal, Canada, as described in the following publication: Tam A, Dansereau C, Badhwar A, Orban P, Belleville S, Chertkow H, Dagher A, Hanganu A, Monchi O, Rosa-Neto P, Shmuel A, Wang S, Breitner J and Bellec P for the Alzheimer’s Disease Neuroimaging Initiative (2015). Common effects of amnestic mild cognitive impairment on resting-state connectivity across four independent studies. *Front. Aging Neurosci.* 7:242. doi: 10.3389/fnagi.2015.00242. It includes group brain parcellations generated from resting-state functional magnetic resonance images for 99 cognitively normal elderly persons and 129 patients with mild cognitive impairment. The brain parcellations have been generated using a method called bootstrap analysis of stable clusters (BASC, Bellec et al., 2010) and eight resolutions (4, 6, 12, 22, 33, 65, 111, and 208 parcels) were selected using a data-driven method called MSTEPS (Bellec, 2013). This work also includes parcellations that contain regions-of-interest (ROIs) that are spatially connected and span only one hemisphere at 8 resolutions (10, 17, 30, 51, 77, 137, 199, and 322 total ROIs). Labels based on typical resting-state networks, and their decomposition into subnetworks or regions, are proposed for all brain parcels. This release also includes unthresholded maps of connectivity differences (t-maps) between patients and controls for four seeds/regions of interest (superior medial frontal cortex, dorsomedial prefrontal cortex, striatum, middle temporal lobe). These seeds were selected because they were associated with the largest number of significant differences between the two populations, as reported in (Tam et al., 2015).

This release more specifically contains the following files:

- README.md: a markdown (text) description of the release.
- brain_parcellation_mcinet_basc_(sym,asym)_(4,6,12,22,33,65,111,208)clusters.(mnc,nii).gz: 3D volumes (either in .mnc or .nii format) at 3 mm isotropic resolution, in the symmetric (symn) and asymmetric (asym) MNI non-linear 2009a space (http://www.bic.mni.mcgill.ca/ServicesAtlases/ICBM152NLin2009), at multiple resolutions of bihemispheric clusters (4, 6, 12, 22, 33, 65, 111, 208).
- labels_mcinet_(sym,asym)_ (12,22,33,65,111,208)clusters.csv: spreadsheets containing labels for each cluster for resolutions containing 12 or more clusters
- brain_parcellation_mcinet_basc_(sym,asym)_ (10,17,30,51,77,137,199,322)rois.(mnc,nii).gz: 3D volumes (either in .mnc or .nii format) at 3 mm isotropic resolution, in the MNI non-linear 2009a space, at multiple resolutions of ROIs (10, 17, 30, 51, 77, 137, 199, and 322 total ROIs), that span only one hemisphere.
- labels_mcinet_(sym,asym)_ (30,51,77,137,199,322)rois.csv: spreadsheets containing labels for each ROI for resolutions containing 30 or more ROIs
- ttest_ctrlvsmci_seed(2,9,12,28).(mnc,nii).gz: 3D volumes (either in .mnc or .nii format) displaying functional connectivity differences (uncorrected t-tests) between patients with mild cognitive impairment and cognitively normal elderly, for 4 different seeds/regions of interest i.e. striatum (seed #2), dorsomedial prefrontal cortex (#9), middle temporal lobe (#12), superior medial frontal cortex (#28); cluster numbers (2, 9, 12, 28) are taken from the parcellation containing 33 clusters.

## Experimental Design, Materials and Methods

### Participants

We pooled resting-state functional magnetic resonance imaging (fMRI) data from four independent studies: the Alzheimer’s Disease Neuroimaging Initiative 2 (ADNI2) sample, two samples from the Centre de recherche de l’institut universitaire de gériatrie de Montréal (CRIUGMa and CRIUGMb), and a sample from the Montreal Neurological Institute (MNI) (Wu et al., 2014). All participants gave their written informed consent to engage in these studies, which were approved by the research ethics board of the respective institutions, and included consent for data sharing with collaborators as well as secondary analysis. Ethical approval was also obtained at the site of secondary analysis (CRIUGM).

The ADNI2 data used in the preparation of this article were obtained from the Alzheimer’s Disease Neuroimaging Initiative (ADNI) database (adni.loni.usc.edu). ADNI was launched in 2003 by the National Institute on Aging, the National Institute of Biomedical Imaging and Bioengineering, the Food and Drug Administration, private pharmaceutical companies and non-profit organizations, as a $60 million, 5-year public-private partnership representing efforts of co-investigators from numerous academic institutions and private corporations. ADNI was followed by ADNI-GO and ADNI-2 that included newer techniques. Subjects included in this study were recruited by ADNI-2 from all 13 sites that acquired resting-state fMRI on Philips scanners across North America. For up-to-date information, see http://www.adni-info.org.

The final combined sample included 112 cognitively normal elderly subjects (CN) and 143 patients with mild cognitive impairment (MCI). In the CN group, the mean age was 72.0 (s.d. 7.0) years, and 38.4% were men. Mean age of the MCI subjects was 72.7 (s.d. 7.7) years, and 50.3% were men.

All subjects underwent cognitive testing. Exclusion criteria common to all studies included: Contraindications to MRI, presence or history of axis I psychiatric disorders (e.g. depression, bipolar disorder, schizophrenia), presence or history of neurologic disease with potential impact on cognition (e.g. Parkinson’s disease), and presence or history of substance abuse. CN subjects did not meet criteria for MCI or dementia. Those with MCI had memory complaints, objective cognitive loss (based on neuropsychological testing), but had intact functional abilities and did not meet criteria for dementia.

### Imaging data acquisition

All resting-state fMRI and structural scans were acquired on Philips and Siemens 3T scanners. For more detailed information on the imaging parameters, please refer to Tam et al., 2015.

### Computational environment

All experiments were performed using the NeuroImaging Analysis Kit (NIAK^1^) (Bellec et al., 2011) version 0.12.18, under CentOS version 6.3 with Octave^2^ version 3.8.1 and the Minc toolkit^3^ version 0.3.18. Analyses were executed in parallel on the “Guillimin” supercomputer^4^, using the pipeline system for Octave and Matlab (Bellec et al., 2012), version 1.0.2. The scripts used for processing can be found on Github^5^.

### Pre-processing

Each fMRI dataset was corrected for slice timing; a rigid-body motion was then estimated for each time frame, both within and between runs, as well as between one fMRI run and the T1 scan for each subject (Collins and Evans, 1997). The T1 scan was itself non-linearly co-registered to the Montreal Neurological Institute (MNI) ICBM152 stereotaxic symmetric template (Fonov et al., 2011), using the CIVET pipeline (Ad-Dab’bagh et al., 2006). The rigid-body, fMRI-to-T1 and T1-to-stereotaxic transformations were all combined to resample the fMRI in MNI space at a 3 mm isotropic resolution. To minimize artifacts due to excessive motion, all time frames showing a displacement greater than 0.5 mm were removed (Power et al., 2012). A minimum of 50 unscrubbed volumes per run was required for further analysis (13 CN and 14 MCI were rejected from the original samples). The following nuisance covariates were regressed out from fMRI time series: slow time drifts (basis of discrete cosines with a 0.01 Hz high-pass cut-off), average signals in conservative masks of the white matter and the lateral ventricles as well as the first 3 to 10 principal components (median numbers for ADNI2, CRIUGMa, CRIUGMb, and MNI were 9, 6, 7, and 7 respectively, and accounting for 95% variance) of the six rigid-body motion parameters and their squares (Giove et al., 2009; Lund et al., 2006). The fMRI volumes were finally spatially smoothed with a 6 mm isotropic Gaussian blurring kernel. A more detailed description of the pipeline can be found on the NIAK website^6^ and Github^7^.

### Parcellation of the brain into functional clusters

After pre-processing, we generated functional brain atlases at multiple resolutions. First, to reduce each fMRI dataset into a time x space array with 957 regions, we applied a region-growing algorithm (Bellec et al., 2006). Next, we applied a bootstrap analysis of stable clusters (BASC) to identify clusters that consistently exhibited similar spontaneous BOLD fluctuations in individual subjects, and were spatially stable across subjects (Bellec et al., 2010). The BASC clustering procedure was carried out at a specific number of clusters (called resolution). Using a “multiscale stepwise selection” (MSTEPS) method (Bellec, 2013), we determined a subset of resolutions that provided an accurate summary of the group stability matrices generated over a fine grid of resolutions: 4, 6, 12, 22, 33, 65, 111 and 208. These eight resolutions of brain parcellations (Figure 1), registered to both symmetric and asymmetric MNI templates, have been released on figshare^8^ and Neurovault^9^. These eight resolutions were further processed to generate eight parcellations that contain ROIs that are spatially connected and span only one hemisphere (for an example, see Figure 2). These latter parcellations contain 10, 17, 30, 51, 77, 137, 199, and 322 total ROIs.

**Figure 1.**
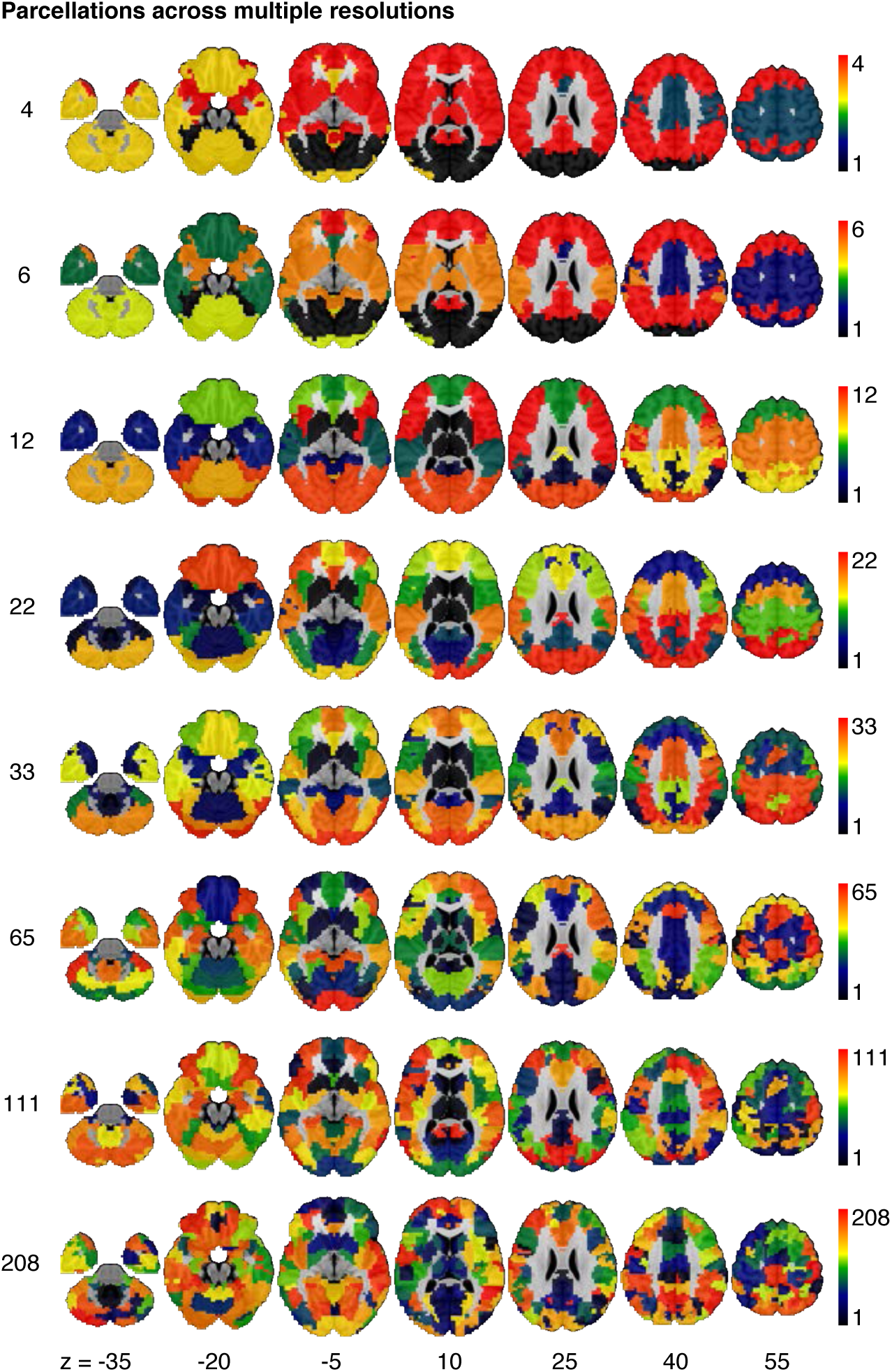
Functional parcellations across resolutions (or number of clusters) selected by MSTEPS (reproduced from Tam et al., 2015).

**Figure 2.**
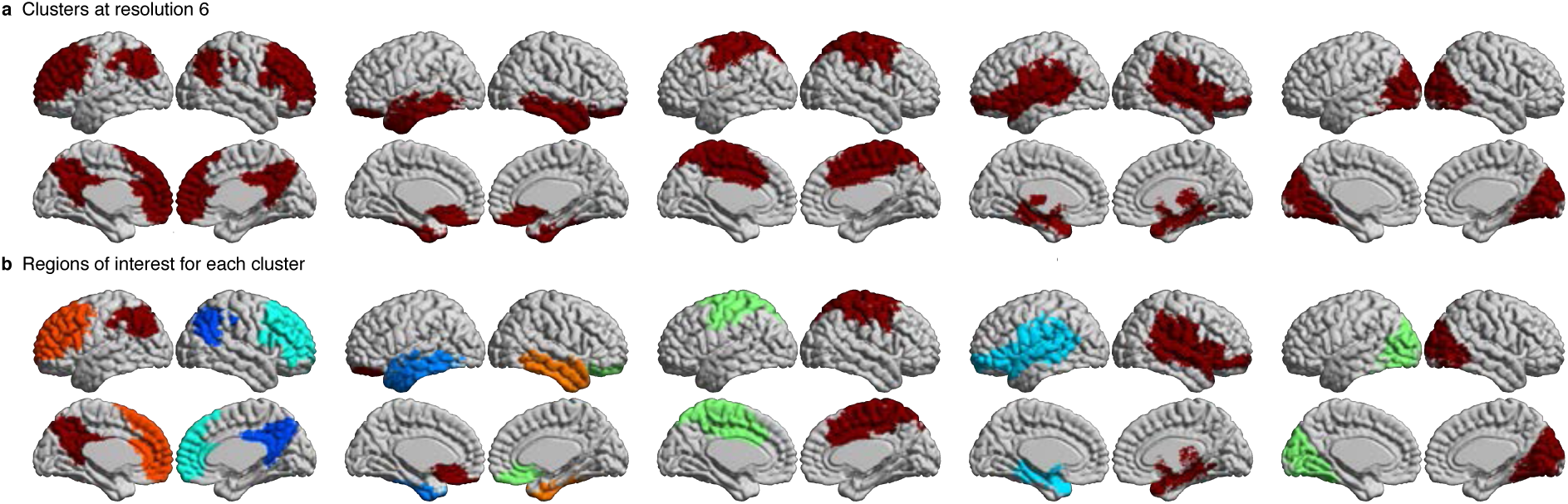
Clusters at resolution 6 (cerebellum not shown) and their respective regions-of-interest. Note how each cluster in (A) is bihemispheric prior to breaking down into multiple spatially constrained regions-of-interest in (B).

We have provided labels for each parcel at every resolution, except for resolutions 4 and 6 due to the merging of networks at those low resolutions. At resolution 4, we observed the sensory-motor network, visual network, a network that resembles the endogenous network (Golland et al., 2008) and a network that merges the cerebellum and the mesolimbic network together. At resolution 6, we observed the visual network, cerebellum, mesolimbic network, sensory-motor network, a network that merges the deep gray matter nuclei with the frontoparietal network, and a network that merges the default mode network with the posterior attention network. For resolution 12, we manually labeled each parcel (deep gray matter nuclei (DGMN), posterior default mode network (pDMN), medial temporal lobe (mTL), ventral temporal lobe (vTL), dorsal temporal lobe (dTL), anterior default mode network (aDMN), orbitofrontal cortex (OFC), posterior attention (pATT), cerebellum (CER), sensory-motor (SM), visual (VIS), and frontoparietal network (FPN)). Then, we decomposed the networks at resolution 12 into smaller subclusters at all higher resolutions (for an example, see Figure 3). Each parcel at higher resolutions was labeled in reference to the parcels at resolution 12, with the following convention: (resolution)_(parcel label)_(#); for example, at resolution (R) 22, the anterior default mode splits into two clusters, which were named “R22_aDMN_1” and “R22_aDMN_2”.

**Figure 3.**
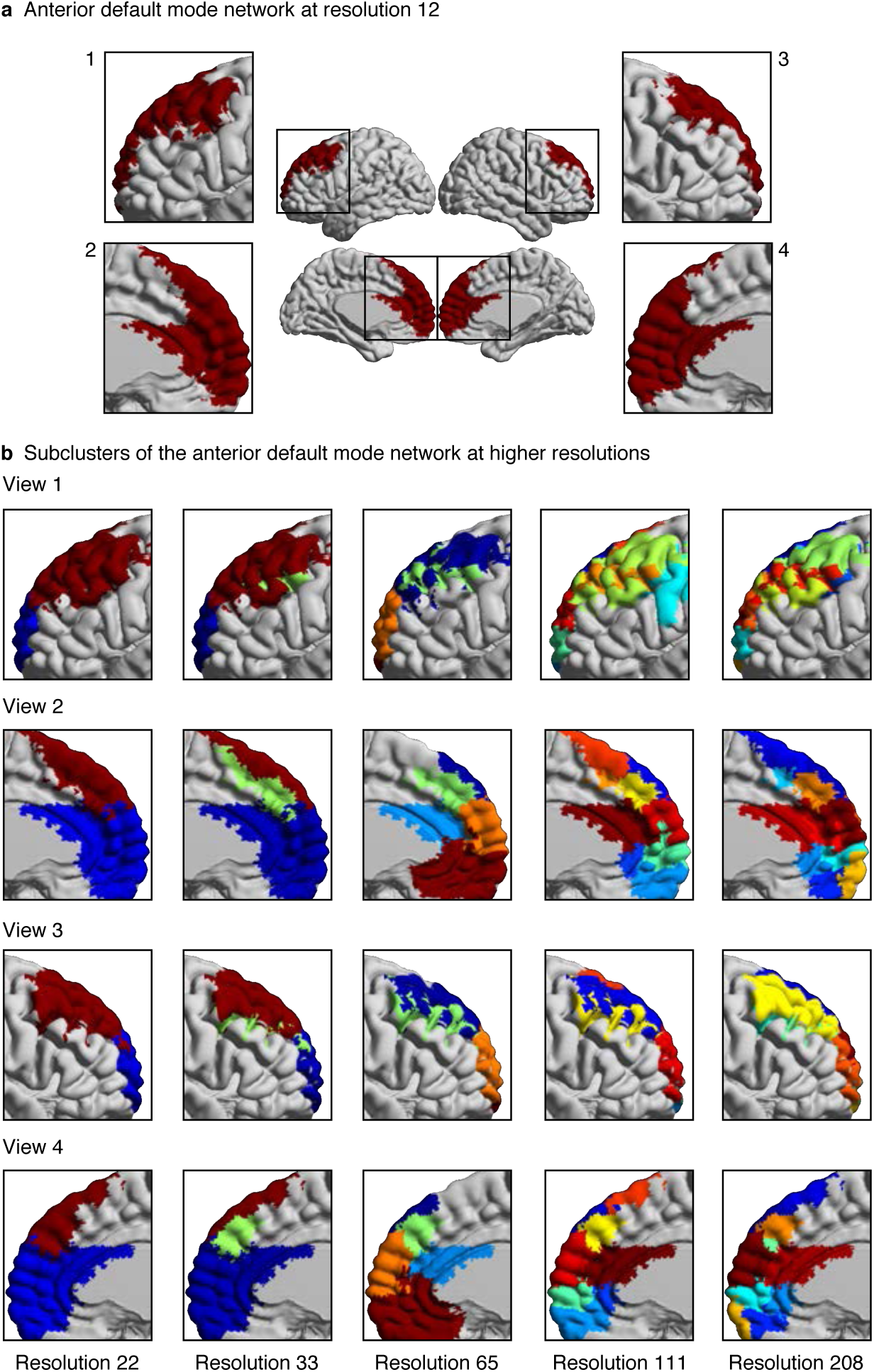
The decomposition of the anterior default mode network into smaller subclusters at higher resolutions in four different views. Resolution 12 was used as a reference for the labeling of subnetworks at higher resolutions.

**Derivation of functional connectomes**

For each resolution *K*, and each pair of distinct clusters, the between-clusters connectivity was measured by the Fisher transform of the Pearson’s correlation between the average time series of the clusters. The within-cluster connectivity was the Fisher transform of the average correlation between time series inside the cluster. An individual connectome was thus a *K x K* matrix.

## Statistical testing

To test for differences between aMCI and CN at a resolution of 33 clusters, we used a general linear model (GLM) for each connection between two parcels (Bellec et al., 2015). The GLM included an intercept, the age and sex of participants, and the average frame displacement of the runs involved in the analysis. The contrast of interest (MCI – CN) was represented by a dummy covariate coding the difference in average connectivity between the two groups. All covariates except the intercept were corrected to a zero mean. The GLM was estimated independently for each scanning protocol. In addition to distinguishing between the CRIUGMa, CRIUGMb, MNI, and ADNI2 samples, ADNI2 was subdivided into five sub-studies based on the use of different Philips scanner models (i.e. Achieva, Gemini, Ingenia, Ingenuity, and Intera). We dropped all subjects scanned with Ingenuity (2 CN, 1 MCI) due to the elimination of all MCI subjects within that site by the scrubbing procedure and its small sample size. We therefore estimated 7 independent GLMs for each protocol (ADNI2-Achieva, ADNI2-Gemini, ADNI2-Ingenia, ADNI2-Intera, CRIUGMa, CRIUGMb, MNI). The estimated effects were combined across all protocols through inverse variance based weighted averaging (Willer et al., 2010). From this analysis, we present uncorrected t-maps illustrating functional connectivity differences between patients and controls for four seeds/regions of interest (superior medial frontal cortex, dorsomedial prefrontal cortex, striatum, middle temporal lobe) (Figure 4). These maps have been released on figshare^8^ and Neurovault^9^. These four seeds were chosen for further analyses because, together, they were associated with 47% of all significant group differences across all brain regions. Briefly, we found that MCI patients exhibited reduced connectivity between default mode network nodes and between areas of the cortico-striatal-thalamic loop. For a more in-depth presentation and discussion of results, please refer to Tam et al, 2015.

**Figure 4.**
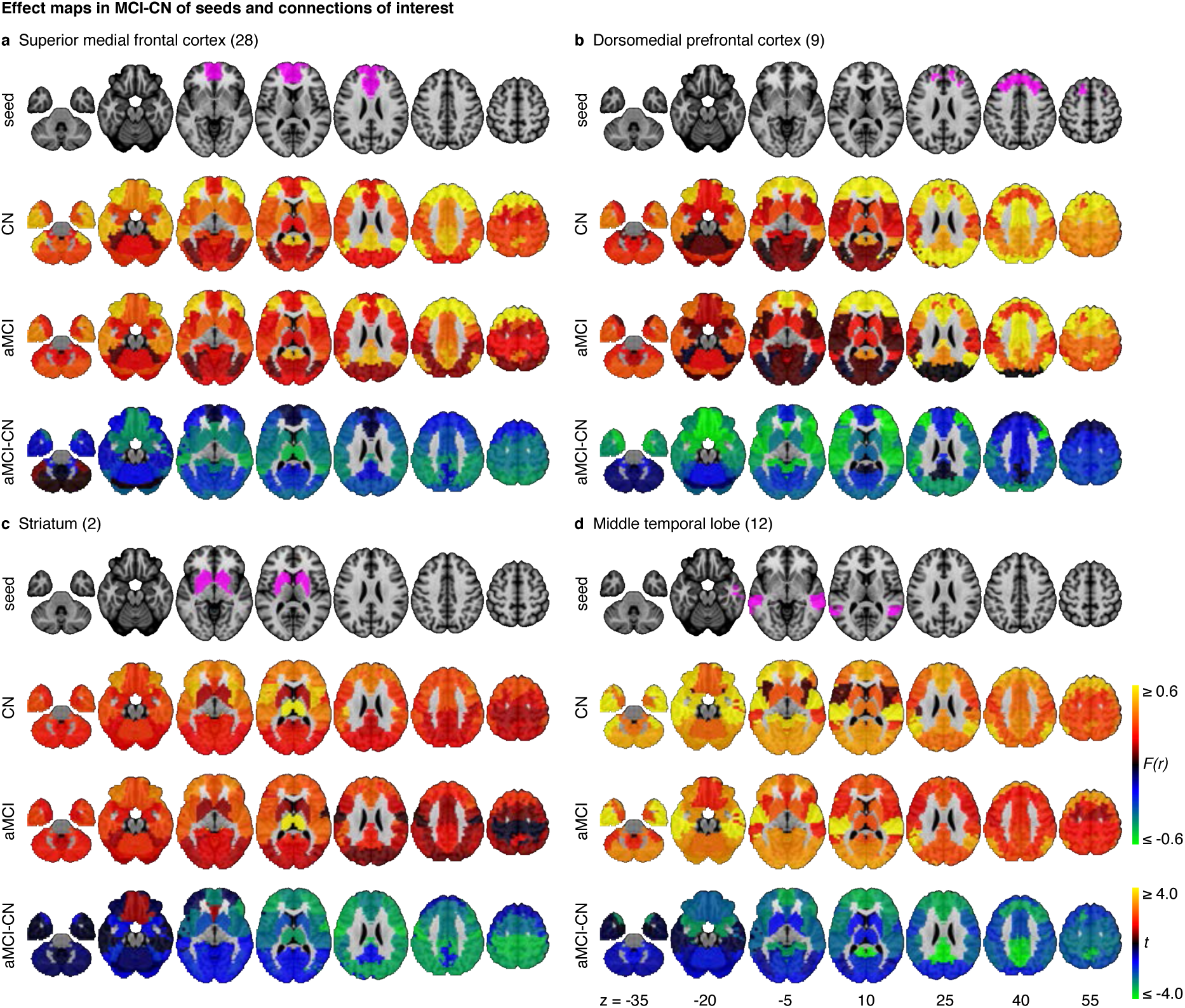
Maps for a selection of four seeds that show effects related to MCI at resolution 33. These effect maps reveal the spatial distribution of the differences in functional connectivity for (A) the superior medial frontal cortex, (B) the dorsomedial prefrontal cortex, (C) striatum, and (D) the middle temporal lobe. For each panel, the top line maps the spatial location of the seed region in magenta, the second and third lines show the connectivity (Fisher-transformed correlation values (F(r)) between the designated seed region and the rest of the brain in CN and MCI respectively, and the fourth line shows a difference map between MCI and CN (t-test). The numbers in parentheses refer to the numerical IDs of the clusters in the 3D parcellation volume at resolution 33 (adapted from Tam et al., 2015).

## Acknowledgements

We thank Sven Joubert, Isabelle Rouleau, and Sophie Benoit for their contribution in collection of the CRIUGMa dataset. We thank Seqian Wang for his help in curating the ADNI dataset. We thank Tom Nichols for his comments and suggestions. Collection of the CRIUGMb dataset was supported by the Alzheimer’s Society of Canada. Collection of the MNI dataset was supported by Industry Canada/Montreal Neurological Institute Centre of excellence in commercialization and research. This work was supported by CIHR (133359) and NSERC (436141-2013). The computational resources used to perform the data analysis were provided by Compute Canada and CLUMEQ, which is funded in part by NSERC (MRS), FQRNT, and McGill University.

Data collection and sharing for this project was funded by the Alzheimer’s Disease Neuroimaging Initiative (ADNI) (National Institutes of Health Grant U01 AG024904) and DOD ADNI (Department of Defense award number W81XWH-12-2-0012). ADNI is funded by the National Institute on Aging, the National Institute of Biomedical Imaging and Bioengineering, and through generous contributions from the following: AbbVie, Alzheimer’s Association; Alzheimer’s Drug Discovery Foundation; Araclon Biotech; BioClinica, Inc.; Biogen; Bristol-Myers Squibb Company; CereSpir, Inc.; Cogstate; Eisai Inc.; Elan Pharmaceuticals, Inc.; Eli Lilly and Company; EuroImmun; F. Hoffmann-La Roche Ltd and its affiliated company Genentech, Inc.; Fujirebio; GE Healthcare; IXICO Ltd.; Janssen Alzheimer Immunotherapy Research & Development, LLC.; Johnson & Johnson Pharmaceutical Research & Development LLC.; Lumosity; Lundbeck; Merck & Co., Inc.; Meso Scale Diagnostics, LLC.; NeuroRx Research; Neurotrack Technologies; Novartis Pharmaceuticals Corporation; Pfizer Inc.; Piramal Imaging; Servier; Takeda Pharmaceutical Company; and Transition Therapeutics. The Canadian Institutes of Health Research is providing funds to support ADNI clinical sites in Canada. Private sector contributions are facilitated by the Foundation for the National Institutes of Health (http://www.fnih.org). The grantee organization is the Northern California Institute for Research and Education, and the study is coordinated by the Alzheimer’s Therapeutic Research Institute at the University of Southern California. ADNI data are disseminated by the Laboratory for Neuro Imaging at the University of Southern California.

## Footnotes

1 http://simexp.github.io/niak/

2 https://www.gnu.org/software/octave/

3 http://www.bic.mni.mcgill.ca/ServicesSoftware/ServicesSoftwareMincToolKit

4 http://www.calculquebec.ca/en/resources/compute-servers/guillimin

5 https://github.com/SIMEXP/mcinet

6 http://niak.simexp-lab.org/pipe_preprocessing.html

7 https://github.com/SIMEXP/mcinet/tree/master/preprocess

8 http://dx.doi.org/10.6084/m9.figshare.1480461

9 http://neurovault.org/collections/1003/

